# A transcriptional continuum links viral burden and host-cell biology in human lymphoid tissue

**DOI:** 10.64898/2026.07.07.736525

**Authors:** Jesse F. Mangold, Nadine Schrode, Trinisia Fortune, Aislinn M. Keane, Sanjana Shroff, Sabrina Petri, Benjamin Tweel, Kristin G. Beaumont, Talia H. Swartz

## Abstract

Human immunodeficiency virus (HIV-1) persistence in lymphoid tissue remains a major barrier to cure, yet infected cells are commonly represented using discrete categories that may obscure biologically meaningful heterogeneity. Using an *ex vivo* human tonsil explant model and single-cell RNA sequencing of more than 42,000 T cells, we show that HIV transcription spans a structured continuum and that viral transcriptional burden functions as an organizing axis of host-cell biology. Across the continuum, increasing HIV transcription is associated with coordinated remodeling of immune, inflammatory, metabolic, and redox-associated programs. Lower transcriptional tiers were enriched for innate sensing and inflammasome-associated responses, whereas higher tiers exhibited activation of oxidative phosphorylation and redox-buffering pathways. Antiretroviral therapy preferentially depleted highly transcriptionally active populations while preserving lower and intermediate tiers, resulting in compression rather than elimination of the continuum. Together, these findings provide a quantitative framework for interpreting HIV transcriptional heterogeneity within human lymphoid tissue and suggest that persistent viral activity reflects a spectrum of host-virus states rather than a single infected-cell population. By linking viral transcriptional burden to distinct host-cell programs, this framework may inform future strategies to reduce HIV persistence and its associated inflammatory consequences.

## Introduction

The majority of HIV-1 replication and persistence occurs within secondary lymphoid tissues, including lymph nodes, gut-associated lymphoid tissue (GALT), and spleen, both before and during antiretroviral therapy (ART) [1–2]. Incomplete penetration of antiretroviral drugs into these compartments has been proposed as a contributor to residual viral replication and reservoir persistence despite suppression of plasma viremia [3]. Understanding HIV biology within lymphoid tissue is therefore essential for advancing curative strategies. Human palatine tonsils provide an accessible source of lymphoid tissue that recapitulates many features of lymph node architecture and immune organization [4–7].

Human tonsil explants preserve native lymphoid architecture and support productive HIV replication ex vivo, providing a tractable system for studying viral persistence and host responses within human lymphoid tissue [24–32]. Prior studies have used tissue explants to characterize inducible HIV reservoirs, identify T follicular helper (Tfh) cells as a major reservoir population in tonsillar tissue, and investigate host-pathogen interactions within physiologically relevant tissue environments [33,34]. In a previous study from our laboratory, this system revealed metabolic remodeling of HIV-infected CD4 T cells, including increased oxidative phosphorylation, while myeloid populations exhibited enrichment of inflammasome-associated pathways [36]. These findings support the utility of tissue-based models for investigating HIV biology beyond what is observable in peripheral blood.

Despite major advances in single-cell genomics, HIV infection is still commonly represented using discrete categories such as infected and uninfected. This framework has been highly informative but may obscure biologically meaningful heterogeneity among HIV-expressing cells. As single-cell technologies increasingly resolve variation in viral transcription [41–47], an important question emerges: how should HIV-exposed and HIV-infected cells be organized in a biologically meaningful and quantitative manner? Here, we hypothesized that viral transcriptional burden serves as a quantitative coordinate of host-cell state, rather than simply distinguishing infected from uninfected cells. Accordingly, we modeled HIV RNA abundance as a continuous biological variable rather than a binary indicator of infection status. Using an ex vivo human tonsil explant model combined with single-cell RNA sequencing, we quantified HIV transcription across more than 42,000 T cells and organized this variation into a data-driven transcriptional continuum. This framework revealed a spectrum spanning unexposed cells, exposed HIV RNA-negative cells, and HIV RNA-positive cells across a broad range of transcriptional activity. Across this continuum, host immune, inflammatory, metabolic, and redox-associated programs varied with viral burden, and ART preferentially depleted highly transcriptionally active populations while preserving lower and intermediate transcriptional tiers. Together, these findings establish a quantitative framework for interpreting HIV transcriptional heterogeneity within human lymphoid tissue and provide new insight into host programs associated with persistent viral activity during ART.

## Methods

### Human tonsil tissue procurement

The research complies with all relevant ethical regulations. Human palatine tonsil tissue was obtained from routine tonsillectomy procedures under an Institutional Review Board–approved protocol at the Icahn School of Medicine at Mount Sinai, STUDY-20-00930. Discarded tissue was collected from de-identified donors who consented to the use of surgical material for research. Exclusion criteria included use of pre-exposure prophylaxis (PrEP). Specimens were procured and processed on the day of surgery.

### Ex vivo tonsil explant culture

Tonsil tissue was trimmed to remove necrotic and adipose material and dissected into approximately 8 mm³ explants. Explants were cultured at an air–liquid interface on collagen sponge (SurgiFoam) rafts placed in net wells within 6-well plates, with 4 mL of culture medium per well. Each raft contained nine explants arranged in a 3×3 matrix. Human tonsil explant tissue blocks and cultures are maintained in RPMI 1640 medium containing 15% FBS, 2 mM GlutaMAX (Life Technologies), 2 mM L-glutamine (Corning), 1 mM sodium pyruvate (Corning), 1% minimal essential medium (MEM) nonessential amino acids (Corning), 2.5 ug/ml amphotericin B (HyClone), 50 mg/ml gentamicin sulfate (Corning), and 0.3 mg/ml Timentin (bioWORLD). Explants were rested overnight at 37°C and on day 0 transferred to fresh medium in the absence of timentin,

### Virus constructs and production

293T cells were transfected with NL4-3 or JR-FL plasmid using transfection reagent Polyjet (Signagen) following the manufacturer’s instructions. Forty-eight hours post-transfection, virus particles were concentrated by centrifugation through a 6% Optiprep solution at 54,000 *g* for 2 hours. Quantification of virus particles was performed with p24 ELISA.

### HIV infection and antiretroviral treatment

Explants were infected on day 0 with replication-competent HIV-1 reporter viruses. CXCR4-tropic NL4-3 (NL-CI) virus expressing mCherry and CCR5-tropic JR-FL virus expressing GFP were used. Viral inoculum was dropped directly onto the surface of each tissue block (8 µL per explant; NL4-3: 10 ng p24; JR-FL: 100 ng p24). Uninfected controls received an equivalent volume of vehicle. Combination antiretroviral therapy (ART) was initiated at day 2 post-infection to allow initial establishment of infection. The ART regimen consisted of tenofovir (5 µM, Sigma-Aldrich), emtricitabine (70 nM, Sigma-Aldrich), and raltegravir (20 nM, Sigma-Aldrich), selected to achieve partial suppression of viral replication. ART-containing media was maintained throughout the culture period. Explants were cultured for up to 8 days with sampling at days 2, 5, and 8.

### Tissue processing and flow cytometry

At each interval, explants were gently agitated by pipetting within-well media onto tonsil blocks to encourage the release of cells into the supernatant. Cells were collected, washed with PBS through 40 um sterile filters, resuspended in 1x PBS 2 mM EDTA 0.5% BSA, and then stained with LIVE/DEAD NIR viability dye and an antibody panel for immune phenotyping. Flow cytometric analysis followed a standardized gating strategy: debris exclusion by FSC/SSC, singlet gating, live cell selection, CD45⁺ leukocyte gating, lineage identification (CD3⁺ T cells, CD4 proxy as CD3⁺CD8⁻), reporter expression (mCherry or GFP) was used to identify productively infected cells. Data were acquired on an Attune NxT flow cytometer and analyzed using FlowJo v 10.10. Cell viability, CD45⁺ frequency, and CD3⁺CD4⁺ T cell frequency were quantified relative to total live cells.

### Single-cell RNA sequencing

Cell suspensions in 1x PBS 0.04% BSA from uninfected, uninfected +ART, NL4-3 infected, and NL4-3 infected +ART conditions collected were processed using a droplet-based single-cell RNA sequencing platform (10x Genomics Chromium). Dissociated suspension tonsil cell viability was assessed by Trypan Blue exclusion, and suspensions with >60% viability and minimal debris were advanced for library preparation using 3′ gene expression chemistry and sequenced on an Illumina platform. Single-cell libraries were generated using the Chromium Single Cell 3′ Gene Expression v3 platform (10x Genomics) with an input target of approximately 10,000 cells per sample. Gel bead-in-emulsion (GEM) generation, reverse transcription, cDNA recovery, amplification, and library construction were performed according to the manufacturer’s protocol. Amplified cDNA underwent fragmentation, end repair, A-tailing, adaptor ligation, and sample indexing. Final libraries were quantified by Bioanalyzer (Agilent) and Qubit fluorometric analysis (Thermo Fisher Scientific) and sequenced in paired-end mode on an Illumina NovaSeq instrument to a target depth of 50,000–100,000 reads per cell. Sequencing reads were processed with Cell Ranger (10x Genomics, v3.0) using a custom combined human GRCh38 reference and HIV NL-CI viral sequence reference, enabling simultaneous quantification of host and viral transcripts.

### Quality control, data integration, and normalization

Single-cell datasets from multiple donors and conditions were integrated using batch correction (Harmony) [48]. Downstream analysis was performed in R using Seurat (v4.4.1) [49]. Genes detected in fewer than three cells were excluded. Cells were filtered using standard quality-control thresholds to remove low-complexity droplets, likely doublets, and damaged cells. Quality control metrics were evaluated by examining the relationships among read depth, transcript counts, feature counts, and mitochondrial genes. Ambient RNA contamination was estimated and corrected using SoupX (v1.6.2) prior to downstream analyses. Exclusion criteria included fewer than 1000 or greater than 80,000 total molecules, fewer than 500 detected genes, mitochondrial transcript content >20%, and low transcript complexity as assessed by the ratio of log10(genes) to log10(UMI) <0.8. In addition, genes expressed in fewer than 0.5% of cells were removed from downstream analysis.

### Quantification of HIV RNA expression

HIV transcripts were quantified per cell and treated as a continuous variable. Cells with zero detectable viral RNA were retained. HIV RNA abundance was treated as a continuous quantitative variable and served as the primary explanatory variable for downstream modeling of host transcriptional responses. For visualization and comparative analyses, this continuous variable was discretized into transcriptional tiers as described below.

### Gaussian mixture modeling and tier assignment

To discretize the HIV transcriptional continuum into interpretable groups, we modeled positive HIV RNA values using Gaussian mixture modeling. This approach identifies boundaries that reflect the underlying distribution of positive HIV RNA values rather than imposing arbitrary thresholds or quantile-based cutoffs. The optimal number of components was selected using the Bayesian Information Criterion (BIC), which balances model fit against complexity and reduces the risk of over- or under-partitioning the data. Each cell was assigned to the component with the highest posterior probability, providing a probabilistic framework for classification while preserving the underlying continuous nature of HIV transcription. The resulting six HIV RNA-positive components were combined with unexposed cells and exposed HIV RNA-negative cells to generate an eight-tier discretized representation of the HIV transcriptional continuum, comprising unexposed (tier 0), exposed HIV RNA-negative (tier 1), low-transcription (tiers 2-3), intermediate-transcription (tiers 4-5), and high-transcription burden states (tiers 6-7).

### Differential expression modeling

Differential gene expression for a continuous covariate (HIV expression) was modeled using a hurdle mixed model while adjusting for donor as a random effect and common cell-level confounders (library size and mitochondrial ratio). For each gene, the log₂ fold change per standard deviation increase in HIV RNA was calculated. Statistical significance was assessed using false discovery rate (FDR) correction. Models were fit for both untreated and ART treated.

Differential gene expression for HIV expression tiers as well as ART-treatment effect were modeled with the built-in Seurat FindMarkers function, using MAST and correcting for donor, library size and mitochondrial ratio.

### Pathway and gene set analysis

Functional enrichment analysis was carried out using both gene set enrichment analysis (GSEA) and over-representation analysis (ORA) implemented in the clusterProfiler R package in conjunction with Reactome pathway annotations [50].

### Statistical analysis

Longitudinal flow cytometry data were analyzed using linear mixed-effects models with donor as a random effect. Fixed effects included condition, time, and their interaction. P-values were adjusted using the Holm method. All statistical analyses were performed in R v4.4.1. Data visualization was performed using ggplot2.

## Results

Fresh palatine tonsil explants were infected with HIV-1 NL4-3 (mCherry) or JR-FL (GFP) fluorescent reporter viruses and cultured with or without antiretroviral therapy (ART) initiated two days after infection (**Fig. 1A**). Productive infection was readily detected by reporter expression in both viral strains, with NL4-3 exhibiting higher levels of infection than JR-FL (**Fig. 1B**). Longitudinal flow cytometric analysis demonstrated accumulation of reporter-positive CD4⁺ T cells in untreated cultures, whereas ART substantially reduced reporter-positive cells in both infection models (**Fig. 1C, Supplementary Table 1**). Explant viability gradually declined over the culture period but remained comparable between untreated and ART-treated conditions (**Fig. 1D**). The proportion of live CD45⁺ leukocytes remained stable throughout the experiment (**Fig. 1E**), and the frequency of CD3⁺ T cells within the CD45⁺ compartment showed minimal change over time (**Fig. 1F**). NL4-3 infection significantly increased reporter-positive cells and reduced the CD4^+^ T cell compartment, whereas ART initiated after infection markedly suppressed productive infection and partially restored CD4⁺ T cell frequency (**Supplementary Table 2**). These data demonstrate productive HIV infection of human tonsil explants and effective suppression by ART administered after infection was established. Tissue viability and immune cell composition was sufficient for downstream single-cell transcriptomic analyses of NL4-3 exposed tonsil cells.

**Fig. 1.**
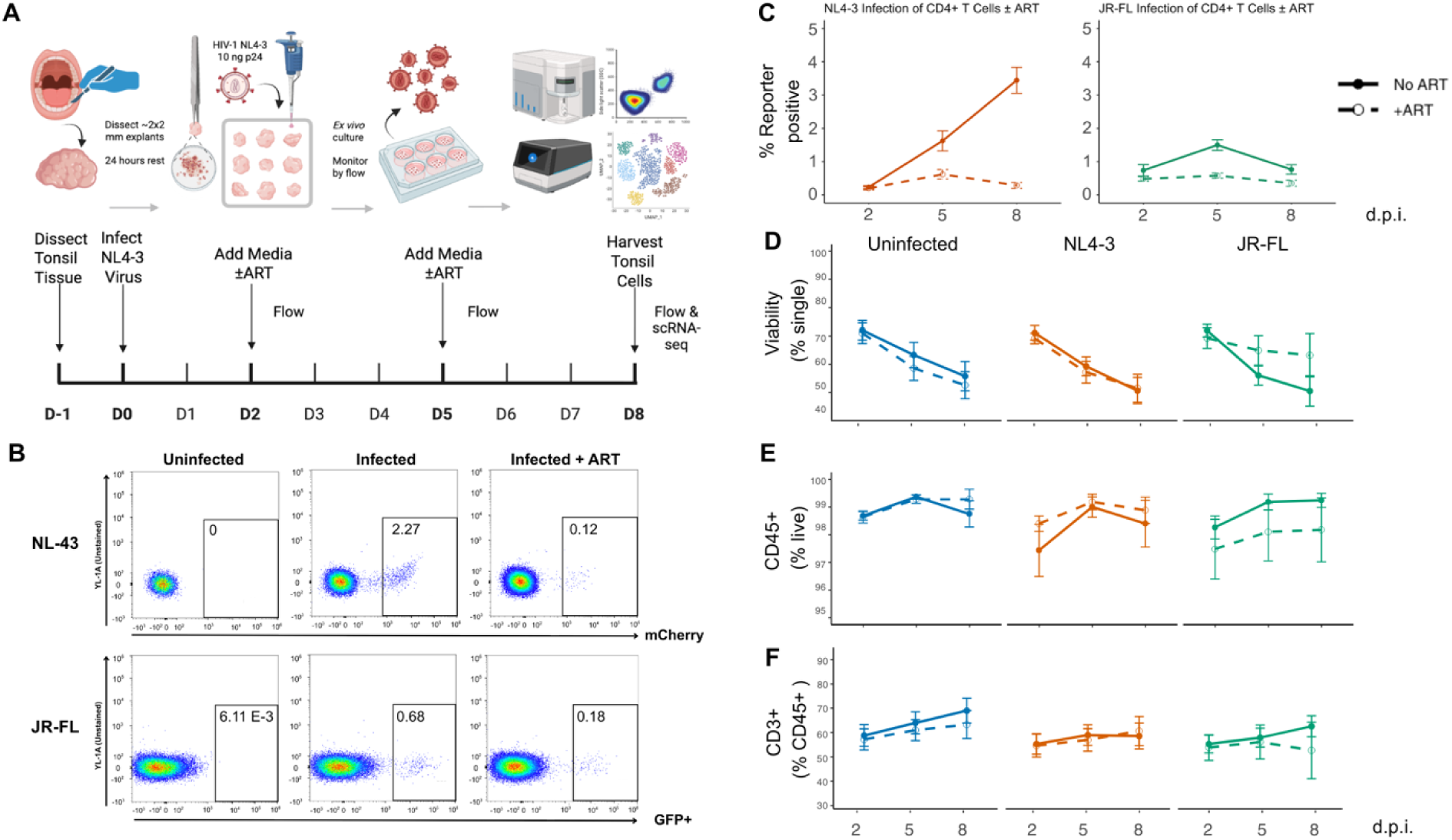
Human tonsil explant model of HIV infection with ART initiated after infection. **A** Experimental workflow. Fresh human palatine tonsil tissue was dissected into approximately 2 × 2 mm explants and cultured at the air–liquid interface. Explants were infected with HIV-1 NL4-3 reporter virus on day 0. Combination antiretroviral therapy (ART) was initiated on day 2 and maintained throughout the remainder of the culture period. Explants were monitored longitudinally by flow cytometry and harvested on day 8 for downstream analyses, including single-cell RNA sequencing. **B** Representative flow cytometry plots showing reporter-positive cells in uninfected controls, infected explants, and infected explants treated with ART. NL4-3 infection was monitored using mCherry expression and JR-FL infection using GFP expression. Values indicate the percentage of reporter-positive cells within the indicated gates. **C** Longitudinal quantification of reporter-positive CD4⁺ T cells following infection with NL4-3 or JR-FL HIV-1 in the presence or absence of ART. Data are shown at 2, 5, and 8 days post-infection (d.p.i.). **D** Percentage of viable single cells over time in uninfected explants and explants infected with NL4-3 or JR-FL HIV-1 cultured with or without ART. **E** Percentage of live CD45⁺ cells over time in uninfected explants and explants infected with NL4-3 or JR-FL HIV-1 cultured with or without ART. **F** Percentage of CD3⁺ T cells within the CD45⁺ compartment over time in uninfected explants and explants infected with NL4-3 or JR-FL HIV-1 cultured with or without ART. Data are presented as mean ± SEM.

Single-cell RNA sequencing of human tonsil explant supernatants generated an atlas of 152,864 cells. Harmony integration across donors demonstrated substantial overlap of cells from individual specimens, indicating minimal donor-driven clustering (**Fig. 2A**). Broad cell-type annotation with the Human Tonsil Atlas identified the major immune and stromal populations expected in secondary lymphoid tissue, including T cells, B cells, NK cells, ILCs, monocytes/macrophages, dendritic cells, granulocytes, epithelial cells, and follicular dendritic cells (**Fig. 2B, Supplementary Table 3**). Because HIV transcription was predominantly detected within the T-cell compartment, subsequent analyses focused on the 42,998 identified T cells. Expression of canonical lineage markers supported annotation of major T-cell populations, including CD4 T cells, CD8 T cells, naïve T cells, cycling T cells, and double-negative T cells (**Fig. 2C**). Detailed annotation of T-cell populations is shown in **Supplementary Fig. 1**. These populations formed a structured T cell manifold that served as the foundation for further interrogation of HIV transcriptional tiers and host responses (**Fig. 2D**). To determine whether viral transcriptional burden could serve as a quantitative organizing axis of host-cell biology, we modeled HIV RNA abundance as a continuous variable and organized cells into data-driven transcriptional tiers (**Fig. 3, Supplementary Table 4**).

**Fig. 2.**
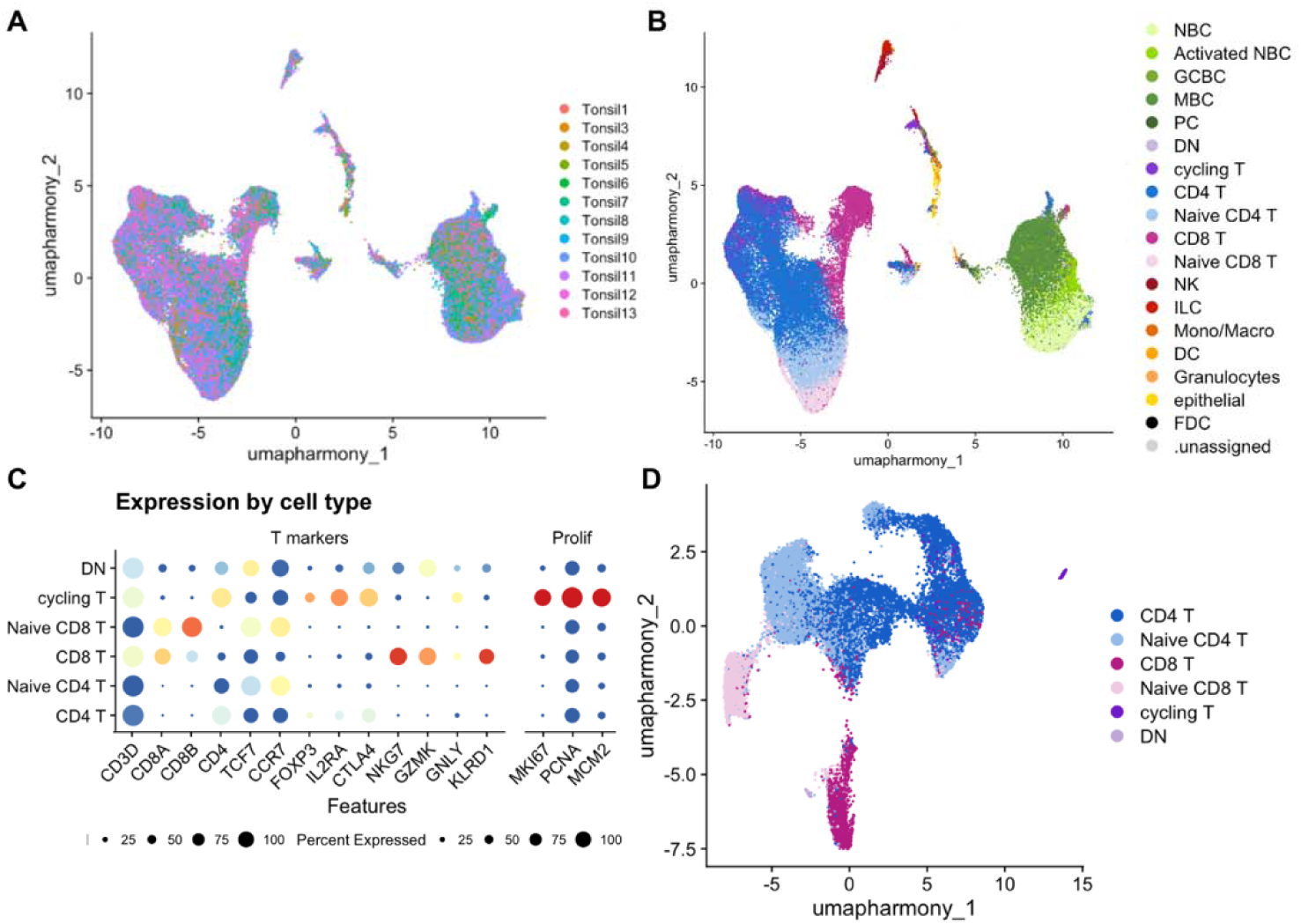
Single-cell atlas of human tonsil explants identifies major immune cell populations and T-cell subsets. **A** Harmony-integrated UMAP of all cells colored by donor. Cells from individual tonsil donors are well mixed across the embedding, indicating successful integration across samples. **B** Harmony-integrated UMAP of all cells colored by broad cell-type annotation with Human Tonsil Atlas. Major immune and stromal populations identified in the tonsil explant model include T cells, B cells, natural killer (NK) cells, innate lymphoid cells (ILCs), monocytes/macrophages, dendritic cells (DCs), granulocytes, epithelial cells, and follicular dendritic cells (FDCs). **C** Dot plot showing expression of representative marker genes used for annotation of T-cell populations. Color indicates average expression and dot size indicates the percentage of cells expressing each gene. **D** UMAP of the T-cell compartment colored by broad T-cell annotation, including CD4 T cells, naïve CD4 T cells, CD8 T cells, naïve CD8 T cells, cycling T cells, and double-negative (DN).

**Fig. 3.**
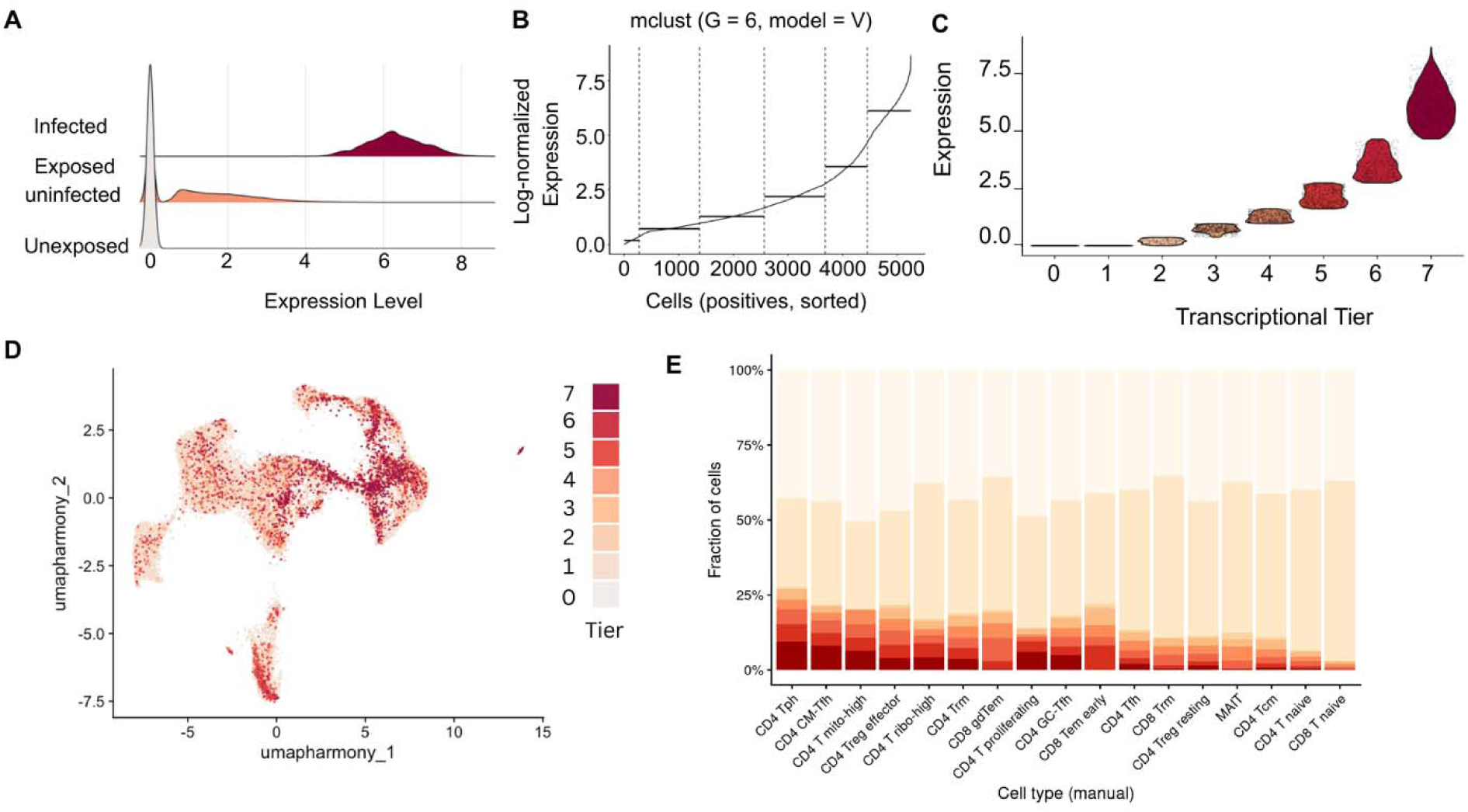
Gaussian mixture modeling provides a discretized representation of the HIV transcriptional continuum. **A** Density distribution of HIV RNA abundance across unexposed cells, exposed HIV RNA-negative cells, and HIV RNA-positive cells. HIV RNA-positive cells spanned a broad range of transcriptional abundance rather than forming a discrete population. **B** HIV RNA-positive cells ordered by HIV RNA abundance (log-normalized expression). Gaussian mixture modeling (mclust; G = 6, model = V) identified six components within the distribution of positive HIV RNA values. The number of components was selected using Bayesian Information Criterion (BIC). **C** Violin plots showing HIV RNA expression across the resulting eight-tier framework, comprising unexposed cells (tier 0), exposed HIV RNA-negative cells (tier 1), and increasing HIV RNA-positive transcriptional tiers (tiers 2–7). **D** Distribution of HIV transcriptional tiers across the integrated Harmony UMAP embedding. Cells are colored according to transcriptional tier. **E** Distribution of HIV transcriptional tiers across manually annotated T-cell subsets in untreated samples. Stacked bar plots show the fraction of cells within each T-cell state assigned to each transcriptional tier.

HIV RNA abundance varied continuously across HIV RNA-positive cells, providing a quantitative basis for resolving host transcriptional programs associated with increasing viral transcription (**Fig. 3A**). Gaussian mixture modeling of positive HIV RNA values identified six data-driven components that, together with unexposed and exposed HIV RNA-negative cells, formed an eight-tier discretized representation of the HIV transcriptional continuum (**Fig. 3B,C**). The optimal number of components was selected using Bayesian Information Criterion (BIC), providing a data-driven discretization of the underlying continuous distribution. Model selection analysis demonstrated that a six-component model provided the optimal balance between fit and complexity (**Supplementary Fig. 2**). These transcriptional tiers were distributed across the tonsillar T cell manifold rather than segregating into isolated clusters, indicating that HIV transcriptional tiers were distributed throughout the tissue landscape (**Fig. 3D**). The continuum framework was reproducible across individual donors, although the relative abundance of specific transcriptional tiers varied between specimens (**Supplementary Fig. 3**). Distribution of transcriptional tiers across T cell subsets revealed substantial heterogeneity in HIV transcriptional burden, with Tph, Tfh-, and Trm-associated populations containing larger fractions of higher transcriptional tiers than naïve T cell populations (**Fig. 3E**). These analyses establish a quantitative framework for organizing host-cell heterogeneity according to viral transcriptional burden. This framework provides a common coordinate system for relating host transcriptional programs to increasing levels of viral transcription across individual cells.

To determine how ART shaped the structure of the HIV transcriptional continuum, we compared the distribution of cells across transcriptional tiers between untreated and ART-treated conditions (**Fig. 4**). ART substantially reshaped the continuum, reducing the frequency of higher HIV transcriptional tiers while preserving lower and intermediate tiers (**Fig. 4A**). Quantification of tier abundance demonstrated enrichment of low-transcription tiers (tiers 1–2) and depletion of higher transcriptional tiers (tiers 3–7), with the strongest reductions observed in tiers 6 and 7 (**Fig. 4C**). This redistribution was observed across multiple T cell subsets (**Fig. 4B**). Examination of cellular composition within each transcriptional tier revealed that higher HIV transcriptional tiers were enriched for activated CD4 T cell populations, including Tfh-, Tph-, and Trm-associated subsets, whereas lower transcriptional tiers contained a larger proportion of naïve and memory populations (**Fig. 4D**). Together, these analyses demonstrate that ART compresses the HIV transcriptional landscape by preferentially depleting high-transcription states while preserving lower and intermediate states.

**Fig. 4.**
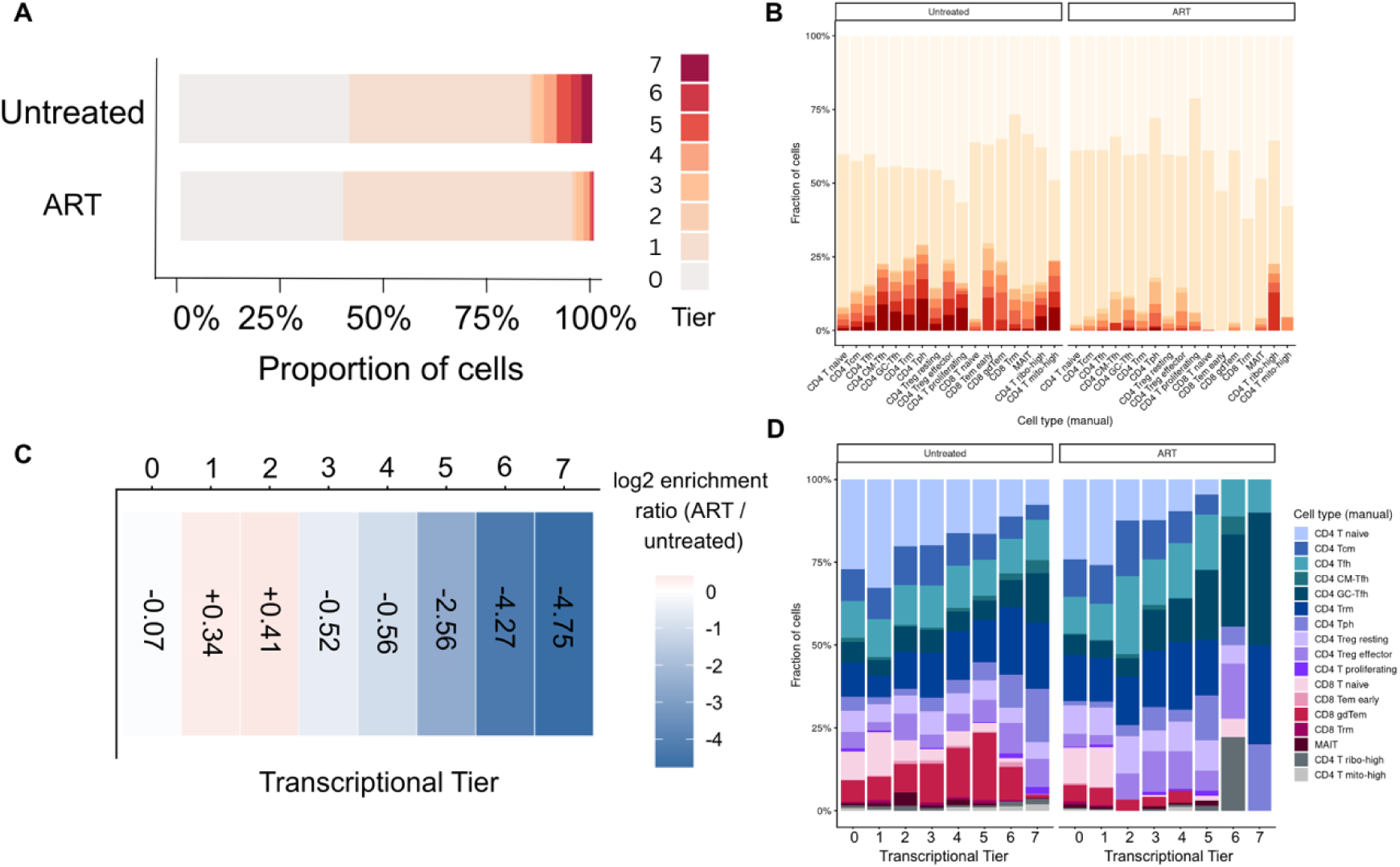
ART reshapes the distribution of HIV transcriptional tiers across the continuum. **A** Distribution of cells across HIV transcriptional tiers under untreated and ART-treated conditions. Stacked bars show the proportion of cells assigned to each transcriptional tier, with cells colored according to transcriptional tier. **B** Distribution of HIV transcriptional tiers across manually annotated T-cell subsets under untreated and ART-treated conditions. Stacked bar plots show the fraction of cells within each subset assigned to each transcriptional tier. **C** Log2 enrichment ratio of cells in each transcriptional tier under ART-treated relative to untreated conditions. Positive values indicate relative enrichment following ART treatment, whereas negative values indicate relative depletion. **D** Cellular composition of each HIV transcriptional tier under untreated and ART-treated conditions. Stacked bar plots show the fraction of cells contributed by each manually annotated T-cell subset within each transcriptional tier.

Using this quantitative framework, we next examined how host transcriptional programs varied along the continuum and how ART reshaped these relationships (**Fig. 5**). Under untreated conditions, increasing HIV burden was associated with transcriptional remodeling, including both negatively associated genes linked to homeostatic, translational, and chromatin-associated programs and positively associated genes linked to activation, signaling, and stress-associated pathways (**Fig. 5A-C**). In contrast, ART-treated cells exhibited markedly reduced effect sizes and fewer strongly associated genes across the continuum (**Fig. 5D**). Despite this attenuation, structured transcriptional programs remained detectable under ART treatment, such as the negative association with *KLF2* and the emergence of a positive association with *IKZF3*. Despite this attenuation, negatively and positively associated gene sets continued to exhibit ordered patterns across transcriptional tiers (**Fig. 5E,F**). These findings support the conclusion that ART compresses HIV-associated host transcriptional remodeling but does not eliminate transcriptional programs associated with increasing HIV burden.

**Fig. 5.**
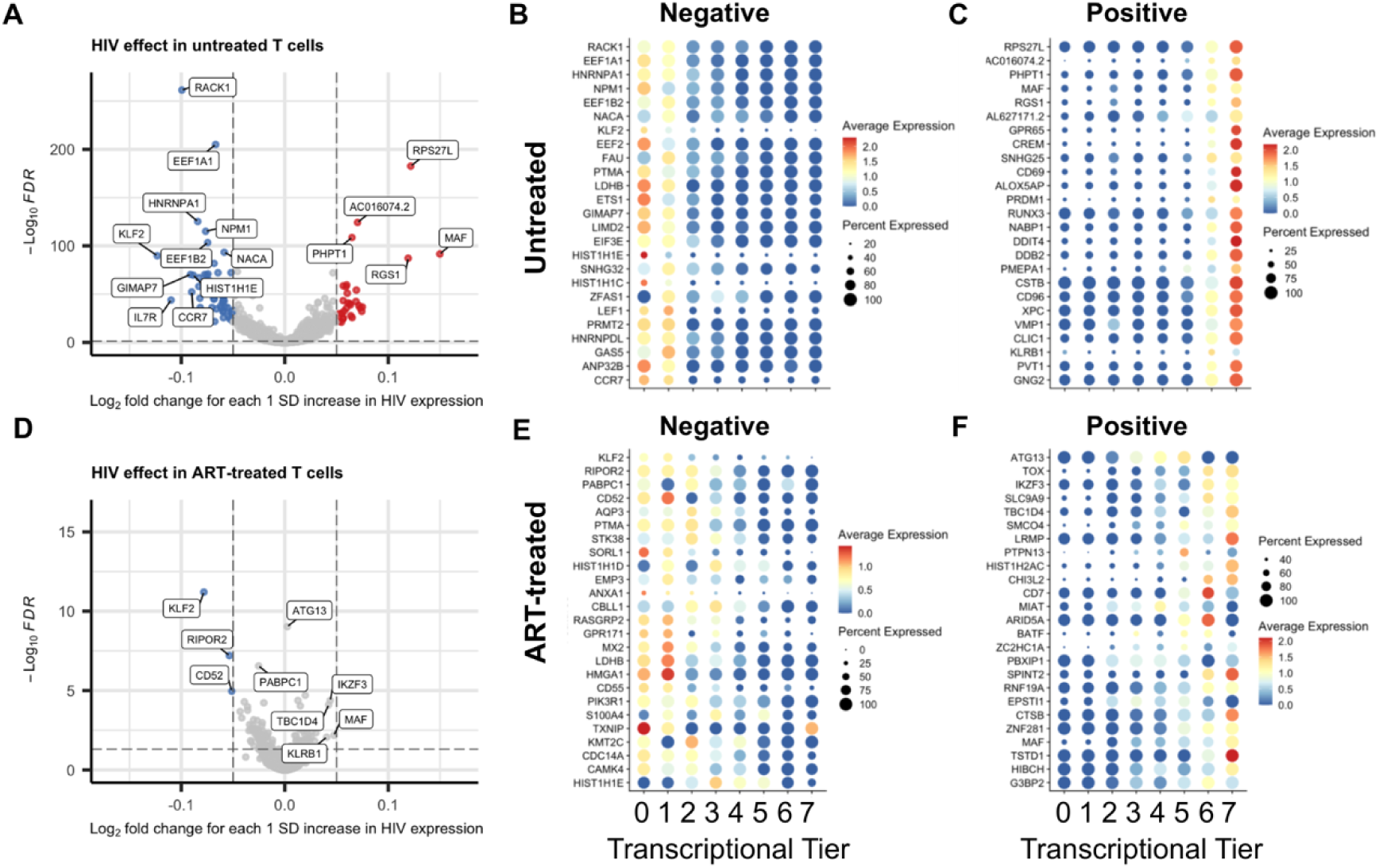
ART attenuates host transcriptional remodeling associated with increasing HIV burden. Differential expression analyses were performed using HIV transcriptional burden as a continuous variable within untreated and ART-treated T cells. Volcano plots show genes associated with increasing HIV expression, with positive log2 fold-change values indicating genes positively associated with increasing HIV burden and negative values indicating genes negatively associated with increasing HIV burden. Selected genes are labeled. Dot plots show representative genes across HIV transcriptional tiers, with color indicating average expression and dot size indicating the percentage of cells expressing each gene. **A** Differential expression associated with increasing HIV burden in untreated T cells. **B** Representative genes negatively associated with increasing HIV burden in untreated T cells. **C** Representative genes positively associated with increasing HIV burden in untreated T cells. **D** Differential expression associated with increasing HIV burden in ART-treated T cells. **E** Representative genes negatively associated with increasing HIV burden in ART-treated T cells. **F** Representative genes positively associated with increasing HIV burden in ART-treated T cells.

To determine whether the biological programs identified across Gaussian mixture-derived transcriptional tiers depended on the discretization strategy, we performed an independent trajectory analysis using pseudotime inferred from the single-cell transcriptomic data (**Fig. 6**). HIV RNA abundance varied systematically along the inferred trajectory (**Fig. 6A**). Representative genes associated with interferon signaling, innate immune activation, and viral sensing exhibited similar graded expression patterns across pseudotime (**Fig. 6B,C**). These findings indicate that the major host transcriptional programs identified using the transcriptional continuum are recapitulated by an independent trajectory inference approach, supporting the robustness of the analytical framework.

**Fig. 6.**
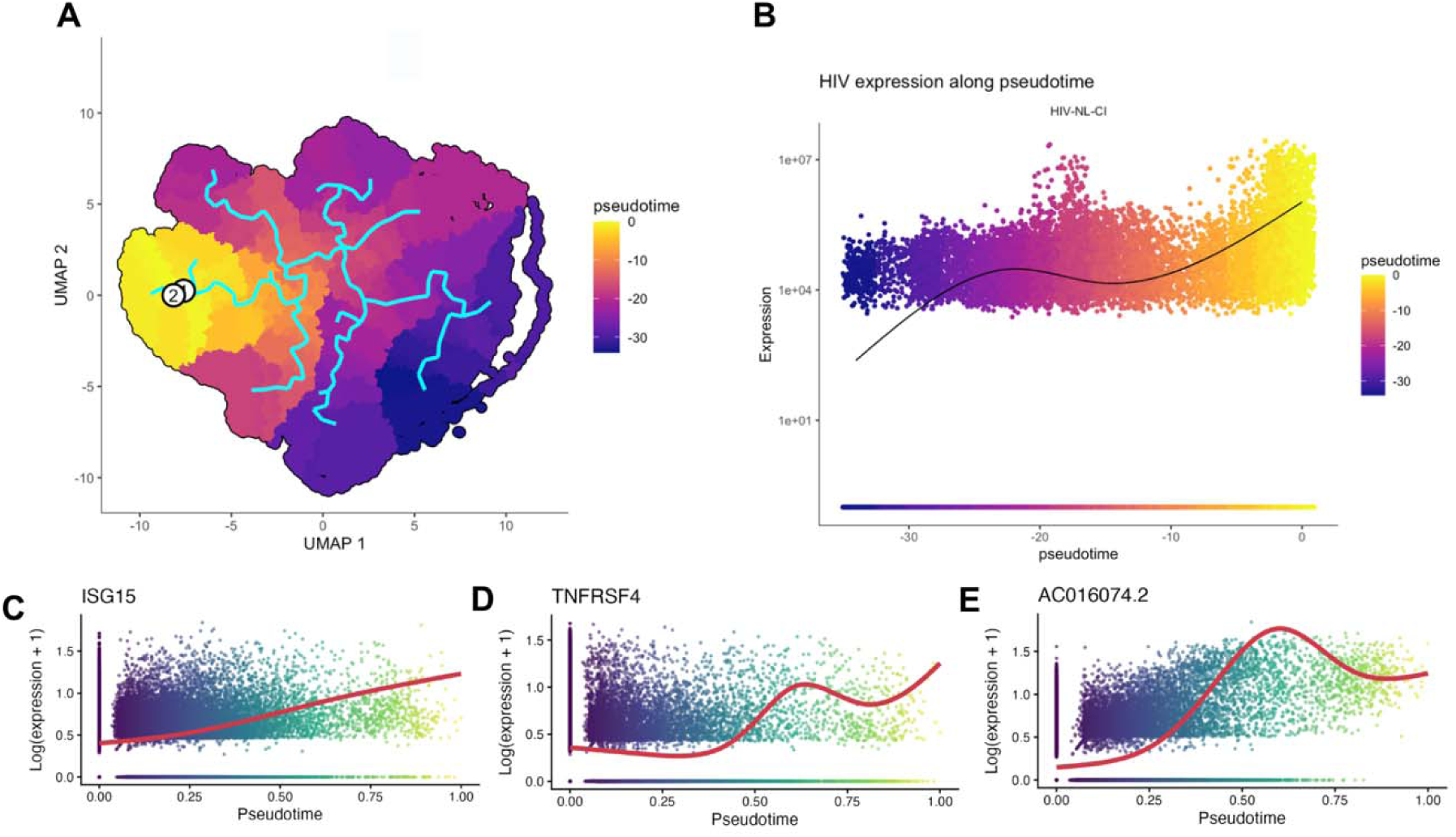
Independent trajectory inference recapitulates host transcriptional programs identified by the HIV transcriptional continuum. **A** UMAP of CD4^⁺^ T cells colored by inferred pseudotime. Cyan lines indicate the inferred trajectory and branch structure, with the circled **1** denoting the inferred starting state. Cells are colored by pseudotime. **B** HIV RNA expression plotted as a function of inferred pseudotime. Each point represents an individual cell colored by pseudotime. The black line indicates a LOESS fit. **C-E** Expression of representative genes across pseudotime. Each point represents a single cell colored by pseudotime. Red lines indicate LOESS fits. **ISG15** (C), **TNFRSF4** (D), and **AC016074.2** (E) are shown as representative genes exhibiting distinct expression patterns across the inferred trajectory.

We next examined biological programs associated with increasing viral transcriptional burden across the continuum (**Fig. 7**). HIV tier 0 (unexposed) was characterized by homeostatic lymphocyte signaling (**Fig. 7A**). HIV tier 1 (exposed HIV RNA-negative) demonstrated enrichment of *IFITM1* expression (**Fig. 7B**). Notably, lower HIV transcriptional tiers also exhibited enrichment of innate immune and inflammasome-associated programs despite relatively limited viral transcription, suggesting that host sensing pathways may be engaged even at low levels of HIV expression. HIV tier 5 exhibited enrichment of long non-coding RNA *AC016074.2*, and at the pathway level featured engagement of T cell activation, antigen presentation, and interleukin-10 signaling, together with repression of WNT-associated pathways (**Fig. 7C**). HIV tier 7 featured significant positive changes in pathways related to respiratory electron transport, reactive oxygen species detoxification, TP53 metabolic pathways, apoptosis-associated programs, and NF-κB signaling, and featured significant negative changes to epigenetic and chromatin pathways **(Fig. 7D**). Together, these analyses reveal coordinated host transcriptional remodeling across the HIV transcriptional continuum, with a higher HIV transcriptional burden associated with coordinated metabolic, redox, and stress-response programs.

**Fig. 7.**
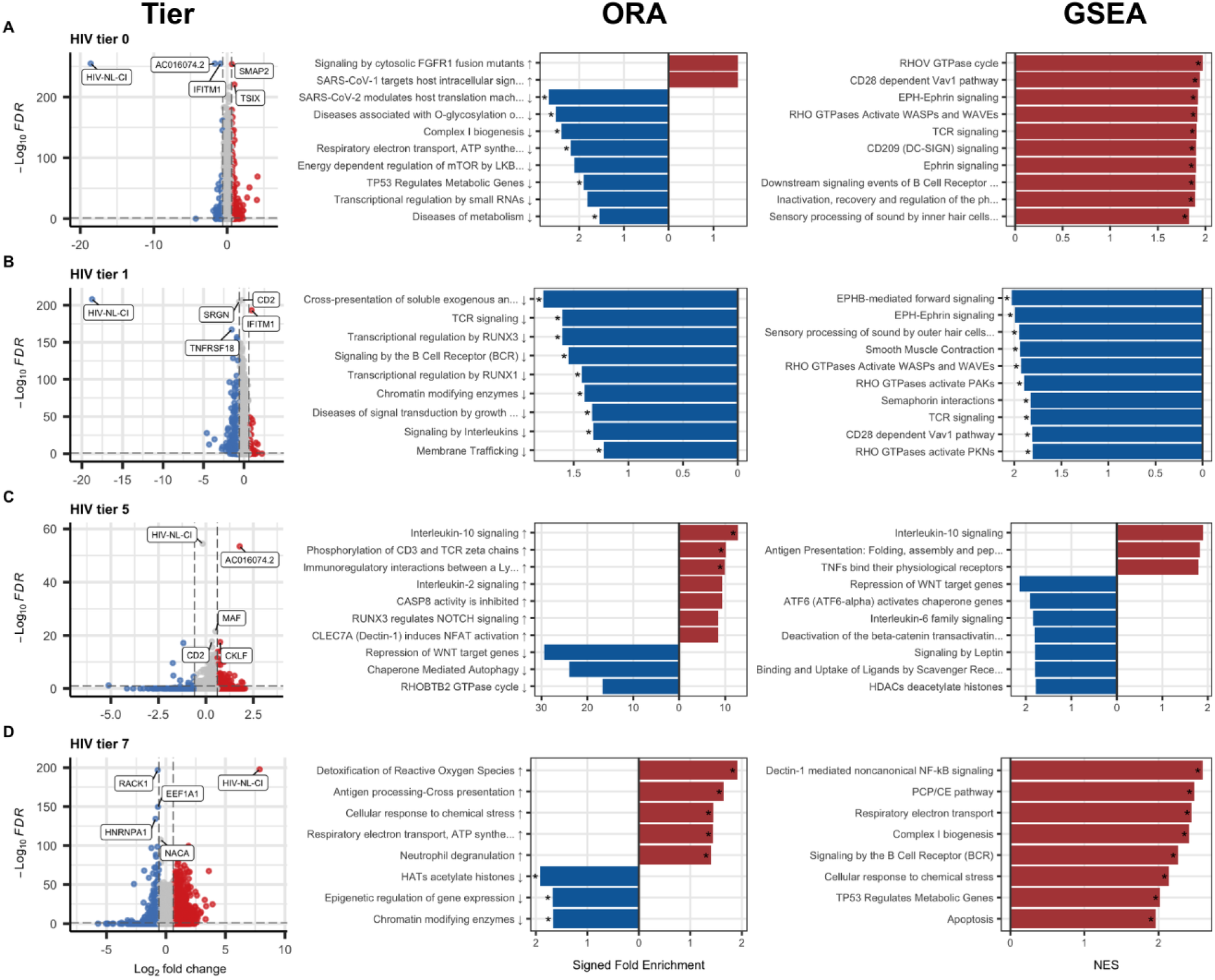
Distinct host transcriptional programs characterize representative positions along the HIV transcriptional continuum. Differential expression analysis was performed for representative HIV transcriptional tiers in untreated T cells, with each tier compared against all remaining tiers. Volcano plots show differential gene expression relative to all other transcriptional tiers, with positive log2 fold-change values indicating genes enriched in the indicated tier and negative values indicating genes relatively depleted from that tier. Selected genes are labeled, including HIV-NL-CI. Ribosomal genes were excluded prior to differential expression and downstream pathway enrichment analyses. Over-representation analysis (ORA) was performed using significantly differentially expressed genes from each tier and is displayed as signed fold enrichment. Gene set enrichment analysis (GSEA) was performed using ranked differential expression statistics and is displayed as normalized enrichment score (NES). Red bars indicate positively enriched pathways and blue bars indicate negatively enriched pathways. Asterisks indicate significantly enriched pathways passing the indicated statistical thresholds. **A** HIV tier 0. **B** HIV tier 1. **C** HIV tier 5. **D** HIV tier 7.

To determine whether HIV positivity was associated with lineage-specific host transcriptional remodeling, we performed differential expression analyses comparing HIV RNA-positive and HIV RNA-negative cells within nine CD4 T cell subsets (**Fig. 8A–I**). Multiple subsets demonstrated reproducible transcriptional remodeling associated with HIV positivity, including recurrent enrichment of genes such as *AC016074.2*, *PMEPA1*, and *RGS16*, and depletion of homeostatic genes such as *IL7R*, *GIMAP7*, and *KLF2*. The magnitude and composition of these transcriptional responses varied across T cell states, indicating both conserved and lineage-specific host programs associated with HIV positivity. CD4 T peripheral helper (Tph) cells exhibited the highest frequency of HIV RNA-positive cells, followed by CD4 CM-Tfh and CD4 mito-high populations, whereas naïve T-cell subsets displayed substantially lower HIV RNA-positive frequencies (**Table 1)**. These findings identify both conserved host responses to HIV positivity and marked differences in susceptibility across tonsillar T cell types.

**Fig. 8.**
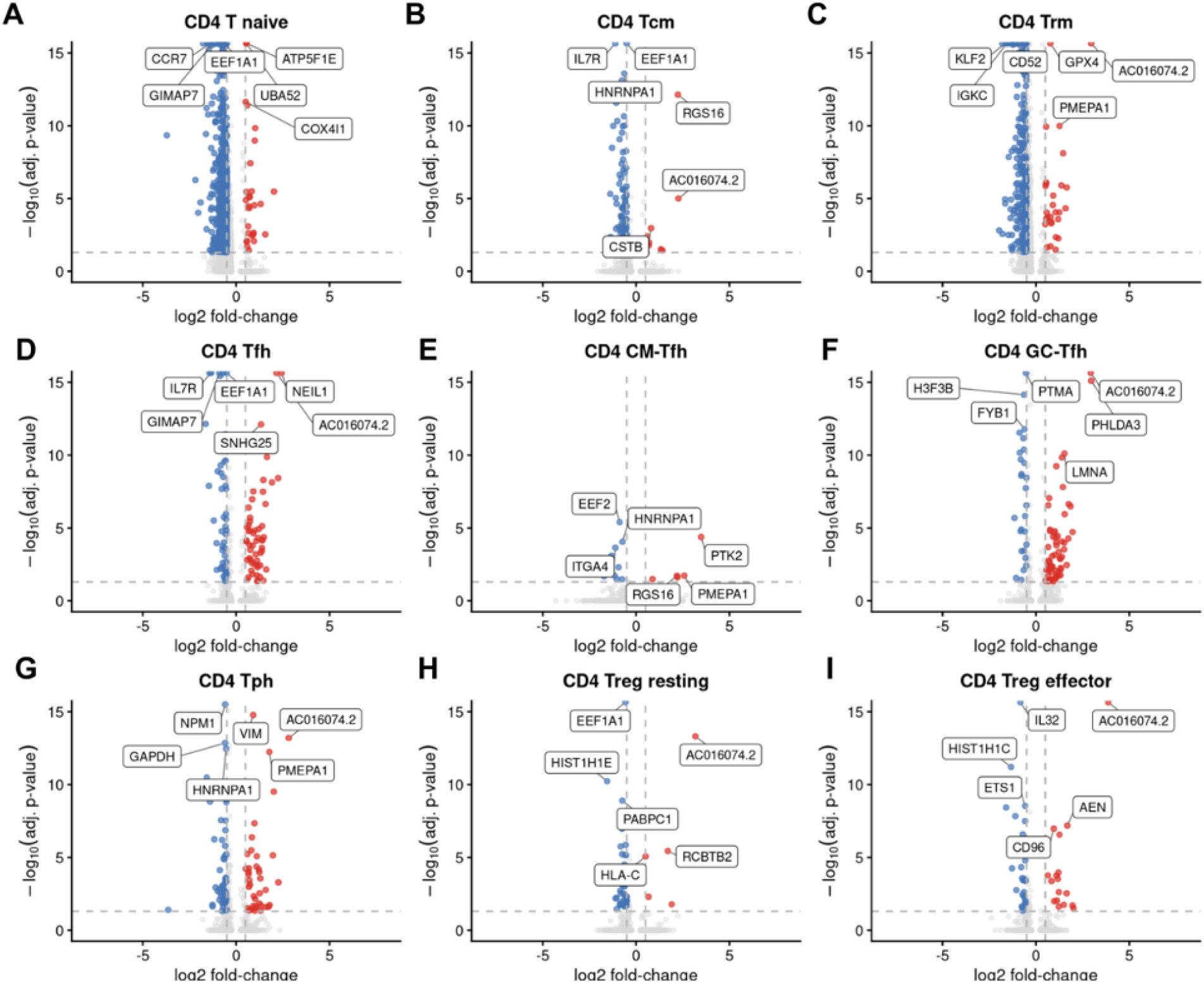
Lineage-specific host transcriptional remodeling associated with HIV positivity in tonsillar T-cell subsets. Differential expression analyses were performed comparing HIV RNA-positive (≥2 UMI cutoff) and HIV RNA-negative cells within individual T-cell subsets. Volcano plots show differential gene expression, with positive log2 fold-change values indicating genes enriched in HIV RNA-positive cells and negative values indicating genes enriched in HIV RNA-negative cells. Selected genes are labeled. Blue points indicate genes negatively associated with HIV positivity and red points indicate genes positively associated with HIV positivity. **(A–I)** Differential expression analyses for individual T-cell subsets: **(A)** CD4 T naive, **(B)** CD4 Tcm, **(C)** CD4 Trm, **(D)** CD4 Tfh, **(E)** CD4 CM-Tfh, **(F)** CD4 GC-Tfh, **(G)** CD4 Tph, **(H)** CD4 Treg resting, and **(I)** CD4 Treg effector.

**Table 1.** Distribution and enrichment of HIV RNA+ cells across tonsillar T cell subsets.

|  |  |  |  |  |
| --- | --- | --- | --- | --- |
| **T cell subset** | **Total Count (n)** | **HIV RNA+ cells (n)** | **Percent of all HIV RNA+ cells** | **Percent HIV RNA+ within** |
| **CD4 Trm** | 4499 | 515 | 17.30% | 11.40% |
| **CD4 T naive** | 11664 | 361 | 12.10% | 3.10% |
| **CD8 gdTem** | 3003 | 337 | 11.30% | 11.20% |
| **CD4 Tfh** | 4784 | 326 | 10.90% | 6.70% |
| **CD4 Tph** | 1442 | 285 | 9.60% | 20.20% |
| **CD4 GC-Tfh** | 2527 | 276 | 9.30% | 10.80% |
| **CD4 Treg effector** | 1722 | 237 | 7.90% | 13.70% |
| **CD4 Tcm** | 4217 | 199 | 6.70% | 4.70% |
| **CD4 Treg resting** | 2823 | 160 | 5.40% | 5.70% |
| **CD4 CM-Tfh** | 389 | 66 | 2.20% | 16.50% |
| **CD8 T naive** | 4285 | 60 | 2.00% | 1.40% |
| **CD4 T mito-high** | 251 | 41 | 1.40% | 16.30% |
| **CD4 T ribo-high** | 322 | 38 | 1.30% | 11.80% |
| **CD4 T prolif.** | 308 | 33 | 1.10% | 10.90% |
| **CD8 Trm** | 4499 | 17 | 0.60% | 6.70% |
| **MAIT** | 366 | 17 | 0.60% | 4.70% |
| **CD8 Tem early** | 145 | 15 | 0.50% | 10.30% |
| Counts and frequencies of HIV RNA+ cells ($\geq 2$ HIV UMIs) across annotated T cell subsets. Percent of all HIV RNA+ cells indicates the contribution of each subset to the total HIV RNA+ pool, whereas Percent HIV RNA+ within indicates the proportion of cells within that subset that are HIV RNA+, a measure of relative enrichment or susceptibility. | | | | |

To further characterize host programs associated with increasing HIV transcriptional burden, we examined mitochondrial respiratory chain, redox-associated, and inflammatory signaling genes across transcriptional tiers under untreated and ART-treated conditions (**Fig. 9**). Genes encoding components of respiratory electron transport chain complexes I, III, IV, and V increased across higher HIV transcriptional tiers under untreated conditions, whereas complex II genes exhibited comparatively limited induction (**Fig. 9A**). Redox-associated genes, including *TXN*, *TXNRD1*, *PRDX1*, *PRDX5*, *TXN2*, and *GLRX*, similarly increased across higher transcriptional tiers, consistent with coordinated induction of antioxidant pathways (**Fig. 9B**). Inflammatory and stress-associated genes, including *NLRP3*, *P2RX7*, *PANX1*, *PYCARD*, *NFKB1*, and *NFKB2*, also varied across transcriptional tiers (**Fig. 9C**). Under ART-treated conditions, many of these programs remained detectable but exhibited altered expression patterns relative to untreated cells. Intriguingly, inflammatory signaling broadly appeared to be more active under ART treatment. Altogether, these analyses identify coordinated mitochondrial, redox-associated, and inflammatory host programs that track with increasing HIV transcriptional burden across the continuum.

**Fig. 9.**
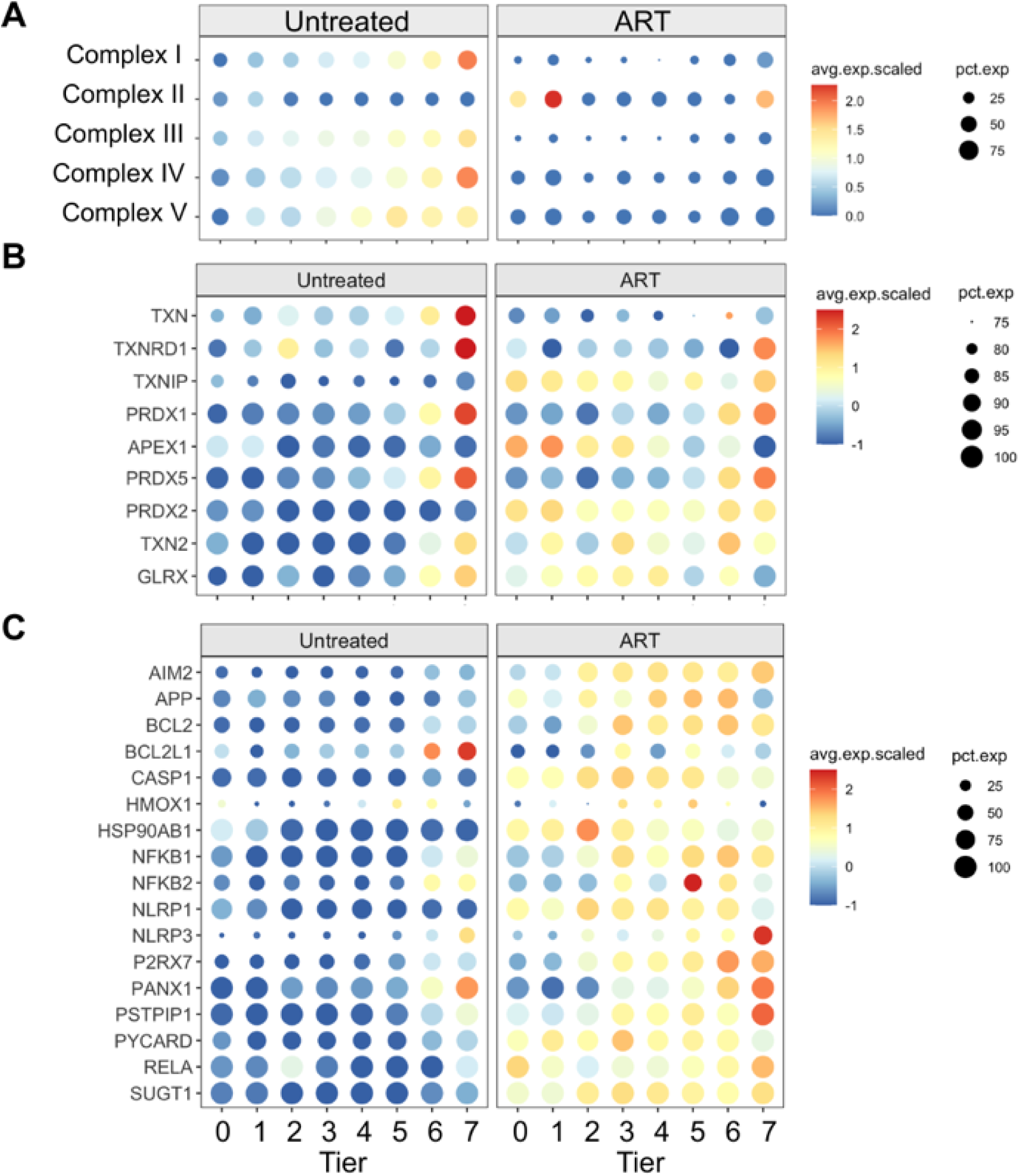
Mitochondrial, redox-associated, and inflammatory programs vary across the HIV transcriptional continuum. Dot plots showing representative mitochondrial respiratory chain, redox-associated, and inflammatory signaling genes across HIV transcriptional tiers under untreated and ART-treated conditions. Color indicates scaled average expression and dot size indicates the percentage of cells expressing each gene. **(A)** Representative genes associated with mitochondrial respiratory electron transport chain complexes I–V across transcriptional tiers under untreated and ART-treated conditions. **(B)** Representative redox-associated genes, including components of the thioredoxin, thioredoxin reductase, glutaredoxin, peroxiredoxin, and oxidative stress response pathways across transcriptional tiers under untreated and ART-treated conditions. **(C)** Representative inflammatory, stress-associated, and inflammasome-related genes across transcriptional tiers under untreated and ART-treated conditions. Transcriptional tiers are shown on the x-axis, with increasing HIV transcriptional burden from tiers 0–7.

## Discussion

In this study, we used an ex vivo human tonsil explant model, combined with single-cell RNA sequencing, to characterize HIV infection at single-cell resolution in cells isolated from their native lymphoid tissue environment. By treating viral transcriptional burden as a continuous quantitative variable rather than a binary indicator of infection status, we identified it as an organizing axis of host-cell biology, linking increased HIV transcription to coordinated remodeling of immune, metabolic, redox, and inflammatory programs. The continuum framework resolves intermediate tiers that are obscured by conventional binary classifications, revealing biologically distinct host programs associated with increasing viral transcriptional burden. Importantly, antiretroviral therapy (ART) does not eliminate this structure but instead redistributes cells across it, selectively depleting high-transcription tiers while preserving lower and intermediate tiers, resulting in compression of the transcriptional continuum (**Fig. 10**). Together, these findings indicate that binary infection categories incompletely capture the biological organization associated with viral transcriptional burden in lymphoid tissue.

**Fig. 10.**
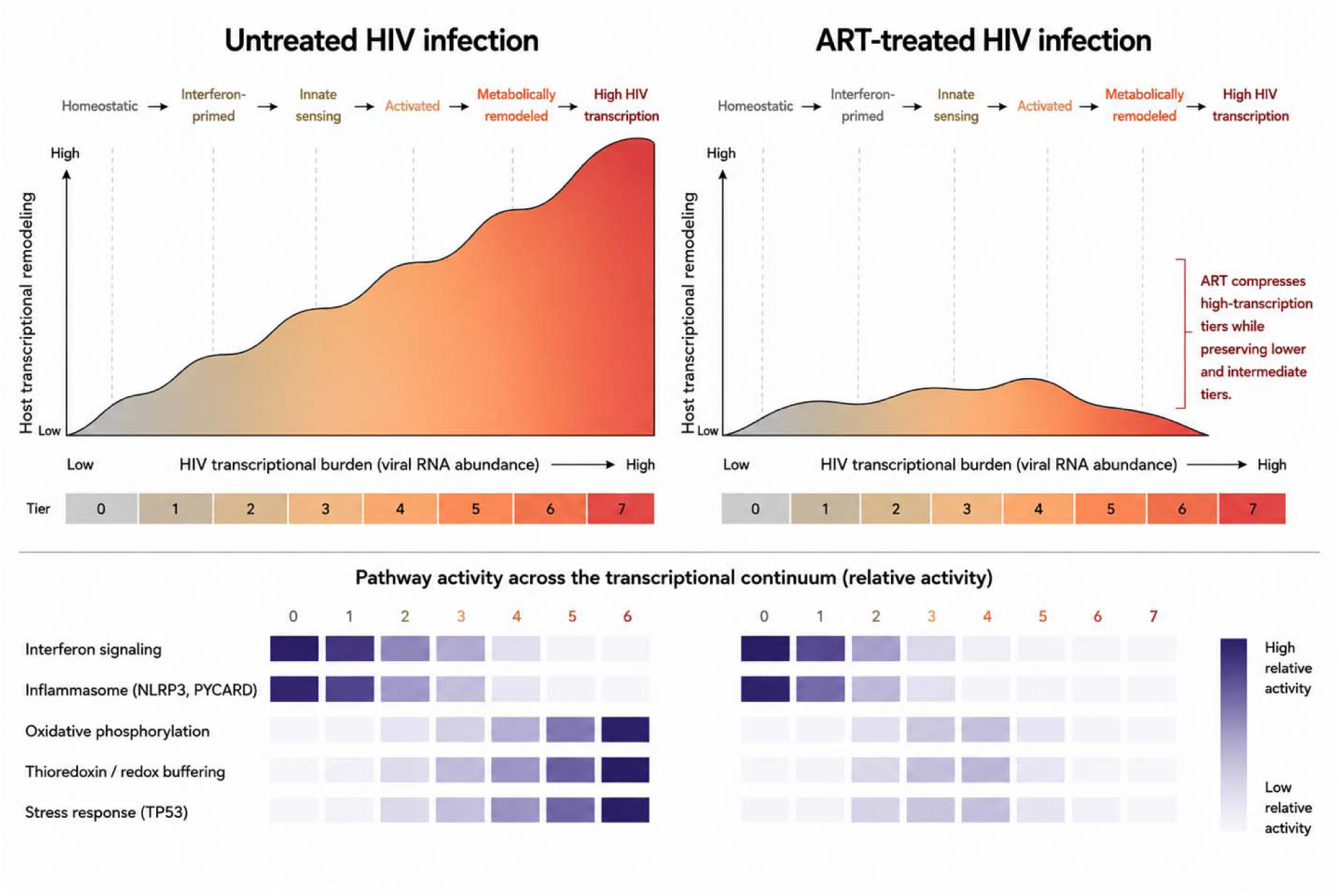
Conceptual model of the HIV transcriptional continuum and its reshaping by antiretroviral therapy. Schematic summary of the transcriptional landscape identified in human tonsil explants. In untreated infection, cells occupy a structured continuum spanning unexposed cells, exposed HIV RNA-negative cells, and progressively higher HIV transcriptional tiers associated with distinct host-cell programs. Antiretroviral therapy preferentially reduces high-transcription states while preserving lower and intermediate transcriptional tiers, thereby compressing the continuum. Representative pathway activity across the continuum is shown schematically beneath each landscape.

Traditional models of HIV infection classify cells as uninfected, infected, or latent using threshold-based measures of viral RNA. These classifications have been invaluable for understanding HIV biology but necessarily reduce a continuous distribution of viral transcription into discrete categories, limiting the ability to resolve graded host-cell responses associated with increasing viral transcriptional burden. Consistent with this view, our analysis demonstrates that HIV RNA abundance spans a continuous distribution without clear separation into discrete transcriptional states, and that intermediate transcriptional tiers are abundant and reproducible across donors. These findings support a model in which host-cell biology varies continuously with viral transcriptional burden, a relationship that is incompletely captured by discrete infection categories. The intermediate tiers identified here should not be interpreted as sequential stages of infection, but rather as data-driven subdivisions of a continuous distribution that facilitate analysis of host transcriptional programs as viral transcriptional burden increases. Within this framework, host responses vary in a coordinated but gene-specific manner across the continuum.

The enrichment of *IFITM1* within tier 1 (exposed HIV RNA-negative) was notable and is consistent with an engagement of anti-viral responses in this population. Interestingly, *IFITM1* has also been linked to inflammatory and epigenetic signatures in HIV-associated aging cohorts, including associations with soluble CD14-related DNA methylation programs [51, 52]. Although our study did not directly assess methylation, these findings raise the possibility that exposure-associated interferon responses in lymphoid tissue may contribute to broader or longer-term inflammatory or epigenetic remodeling in bystander cells lacking detectable HIV RNA.

The observation that lower HIV transcriptional tiers exhibited innate immune and inflammasome-associated signatures despite relatively limited viral transcription suggests that host sensing pathways may be activated early along the HIV transcriptional continuum. This finding is consistent with prior studies demonstrating inflammatory responses during abortive or low-level HIV infection and raises the possibility that innate sensing mechanisms contribute to host transcriptional remodeling even in cells with minimal detectable HIV transcription.

Differential expression across transcriptional tiers and within T cell subsets identified shared top DEGs, including positive association with AC016074.2 and negative association with KLF2. The long non-coding RNA AC016074.2 (*LINC02964*) is previously described in a study of Tat-LNP stimulation of CD4^+^ T cells from PWH on ART describing it as the top hit between p24+ and p24-cells [43]. This molecule may represent a novel HIV-1 responsive regulatory RNA involved in active viral replication. In contrast, recent studies have identified KLF2, a quiescence regulator, as a key transcription factor among HIV infected CD4+ T cells along the path towards latency [53, 54]. Altogether, this continuum framework reveals biology that is consistent with associations previously described in the literature.

Among CD4 T-cell subsets, Tph cells exhibited the highest frequency of HIV transcription, suggesting that peripheral helper-like populations may represent an underappreciated niche for active HIV transcription within lymphoid tissue. Given the emerging role of Tph cells in chronic inflammation and ectopic lymphoid responses, these findings warrant further investigation of their potential contribution to HIV persistence.

A prominent feature of this continuum is the scaling of metabolic programs with HIV transcriptional burden. Expression of mitochondrial respiratory chain gene signatures increases across higher transcriptional tiers, indicating that cells with greater viral RNA levels are associated with enhanced metabolic activity. These findings are consistent with prior work demonstrating that HIV infection is linked to metabolic reprogramming but extend this concept by showing that metabolic programs track the intensity of viral transcription at single-cell resolution. In this context, metabolism appears not only as a permissive factor for infection but as a graded feature of the host–virus interaction.

In parallel, redox-associated pathways, including components of the thioredoxin system, also increase across higher transcriptional tiers. Redox balance is known to regulate transcription factor activity and cellular stress responses and has been implicated in HIV transcriptional control. The observed coupling between redox gene expression and viral RNA levels suggests that oxidative and reductive pathways may help define cellular states along the continuum, potentially influencing both viral transcription and host cell survival.

ART treatment reshapes this transcriptional landscape in a distinct manner. This selective effect indicates that ART does not reset cells to a baseline state but instead preserves a residual transcriptional landscape. Rather than uniformly suppressing all states, ART selectively reduces the abundance of cells in high-transcription tiers while increasing the relative representation of lower and intermediate states. At the level of host gene expression, ART attenuates the association between HIV RNA and metabolic programs, with diminished enrichment of mitochondrial pathways across tiers. In contrast, inflammatory and immune signaling pathways remain detectable across transcriptional tiers and do not exhibit the same degree of attenuation. Together, these observations indicate that ART compresses the transcriptional continuum by limiting high-level viral transcription while preserving a residual activated state.

The persistence of intermediate transcriptional tiers under ART has potential implications for HIV pathogenesis and persistence. These cells may represent a transcriptionally active population that falls below thresholds typically used to define productive infection and may contribute to ongoing immune activation despite effective viral suppression. More broadly, these findings suggest that HIV persistence may involve a spectrum of transcriptionally distinct cellular states with differing biological properties.

This study has several limitations. Although ambient RNA contamination was mitigated using SoupX-based correction, low-level contamination from extracellular viral transcripts cannot be completely excluded. The *ex vivo* tonsil explant model preserves key features of lymphoid tissue architecture but does not fully recapitulate in vivo conditions, including long-term infection dynamics and systemic immune interactions. The use of reporter viruses and short time frames limits direct inference about chronic infection or latency. In addition, the analyses presented here are correlative and do not establish causal relationships between metabolic or redox pathways and HIV transcription. Future studies incorporating perturbation approaches and in vivo validation will be necessary to determine the mechanistic role of these pathways.

Despite these limitations, our findings provide a framework for understanding HIV transcriptional burden as a continuous and structured variable that organizes host-cell biology. By linking viral transcriptional burden to coordinated host programs and demonstrating how ART reshapes this landscape, this work offers a new perspective on the cellular states that support HIV persistence. This continuum-based model may inform future strategies aimed at targeting specific transcriptional or metabolic states to disrupt viral persistence and improve therapeutic outcomes.

Together, these findings establish viral transcriptional burden as a quantitative organizing axis for host-cell biology in HIV-infected lymphoid tissue. By modeling viral transcription as a continuous biological variable, this framework reveals intermediate host-cell programs that are largely obscured by conventional binary classifications of infected and uninfected cells. Increasing viral transcriptional burden is accompanied by coordinated remodeling of host-cell programs, including interferon signaling, metabolic activation, oxidative phosphorylation, redox buffering, stress responses, and inflammatory pathways. ART substantially reshapes this landscape by preferentially depleting high-transcription states while preserving lower and intermediate states, resulting in compression rather than elimination of the continuum. More broadly, these findings suggest that HIV persistence reflects a spectrum of host-virus states distributed across lymphoid tissue, with implications for understanding reservoir biology and the development of host-directed HIV cure strategies.

## Author Contributions

Conceptualization, THS and KGB; methodology, THS, KGB, TF, AMK, SS, SP, and JFM; software, NS; validation, NS, JFM, TF, KGB, and THS; formal analysis, NS, JFM, TF, KGB, and THS; investigation, TF and JFM; resources, BT and THS; data curation, NS; writing—original draft preparation, NS, TF, JFM, KGB, and THS; writing—review and editing, NS, TF, AMK, JFM, SS, SP, BT, KGB, and THS; visualization, NS, TF, AMK, JFM, KGB, and THS; supervision, KGB and THS; project administration, KGB and THS; funding acquisition, JFM and THS.

## Institutional Review Board Statement

The studies involving human participants were conducted in accordance with the Declaration of Helsinki and were reviewed and approved by The Program for the Protection of Human Subjects Institutional Review Board at the Icahn School of Medicine at Mount Sinai under protocol code STUDY-20-00930. The patients/participants provided their written informed consent to participate in this study.

## Informed Consent Statement

Informed consent was obtained from all subjects involved in the study.

## Data Availability Statement

The data supporting the reported results in this study are available upon reasonable request from the corresponding author.

## Acknowledgments

We thank the patients who contributed tonsils to this study. We also thank the BEI Resources and ATCC for providing critical reagents and the Flow Cytometry CoRE at the Icahn School of Medicine at Mount Sinai for assistance with data collection and analysis. We thank members of the Chen lab for thoughtful discussions. This work was supported in part through the computational and data resources and staff expertise provided by Scientific Computing and Data at the Icahn School of Medicine at Mount Sinai and supported by the Clinical and Translational Science Awards (CTSA) grant UL1TR004419 from the National Center for Advancing Translational Sciences. Research reported in this publication was also supported by the Office of Research Infrastructure of the National Institutes of Health under award number S10OD026880 and S10OD030463. The content is solely the responsibility of the authors and does not necessarily represent the official views of the National Institutes of Health. “Research reported in this publication was supported by the Flow Cytometry CoRE (RRID:SCR_027701) at the Icahn School of Medicine at Mount Sinai and by the National Cancer Institute of the National Institutes of Health under award number P30 CA196521. The content is solely the responsibility of the authors and does not necessarily represent the official views of the National Institutes of Health.”

---

**Supplementary Fig. 1.**
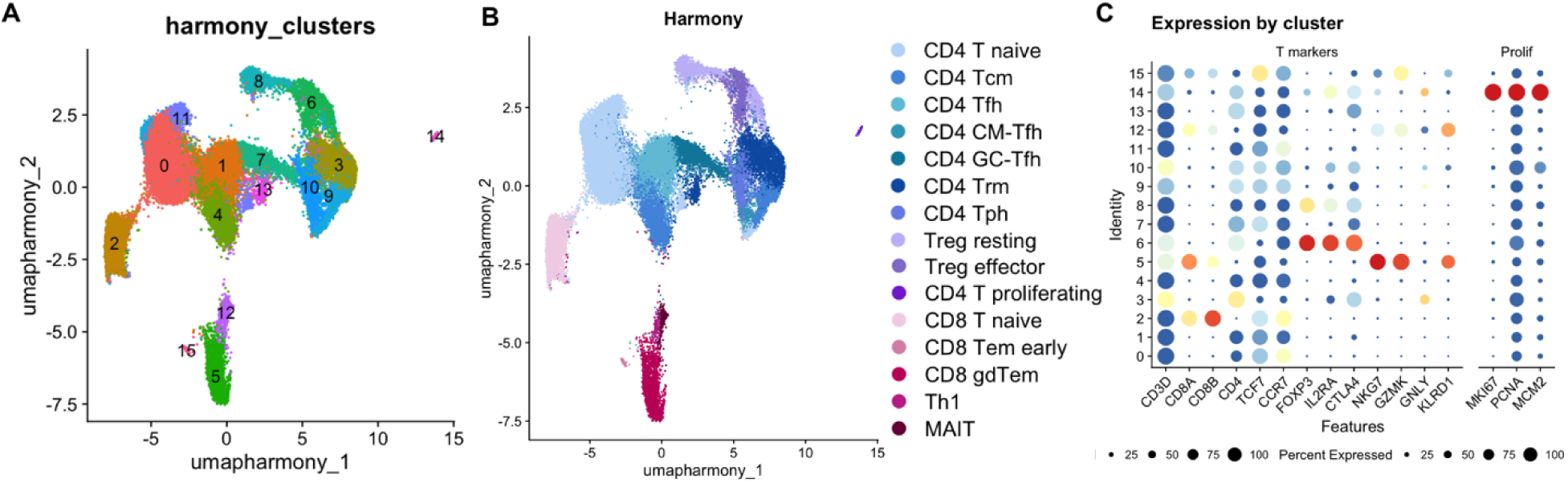
Annotation of the tonsillar T-cell compartment. **A** Harmony-integrated UMAP of T cells colored by unsupervised clustering. Cluster identities were generated following Harmony integration and graph-based clustering. **B** Harmony-integrated UMAP colored by manually curated T-cell annotations. Major T-cell populations identified included CD4 T naive, CD4 Tcm, CD4 Tfh, CD4 CM-Tfh, CD4 GC-Tfh, CD4 Trm, CD4 Tph, resting and effector Treg populations, proliferating CD4 T cells, CD8 T naive, CD8 Tem early, CD8 gdTem, Th1, and MAIT cells. **C** Dot plot showing expression of representative marker genes used for manual annotation of T-cell subsets. Color indicates average expression and dot size indicates the percentage of cells expressing each gene. Canonical lineage markers include *CD3D*, *CD8A*, *CD8B*, *CD4*, *TCF7*, *CCR7*, *FOXP3*, *IL2RA*, *CTLA4*, *NKG7*, *GZMK*, *GNLY*, *KLRD1*, and proliferation-associated genes *MKI67*, *PCNA*, and *MCM2*.

**Supplementary Fig. 2.**
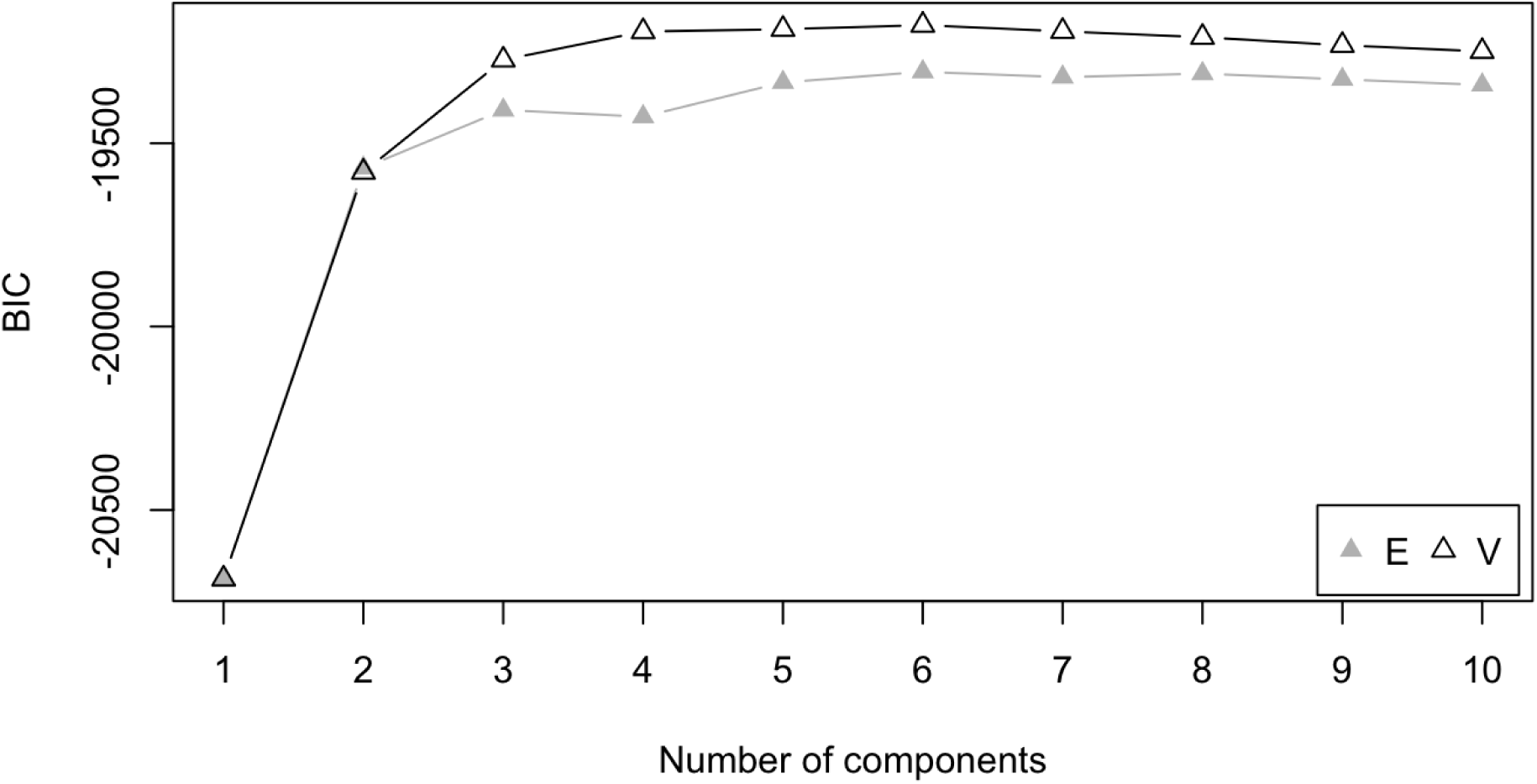
Bayesian Information Criterion (BIC) model selection for Gaussian mixture modeling of HIV RNA-positive cells. Gaussian mixture models with 1–10 components were fit to positive HIV RNA values. The optimal model was selected using Bayesian Information Criterion (BIC), which balances model fit and model complexity. The highest BIC score was observed for the six-component model (G = 6, model = V), which was subsequently used to define HIV transcriptional tiers.

**Supplementary Fig. 3.**
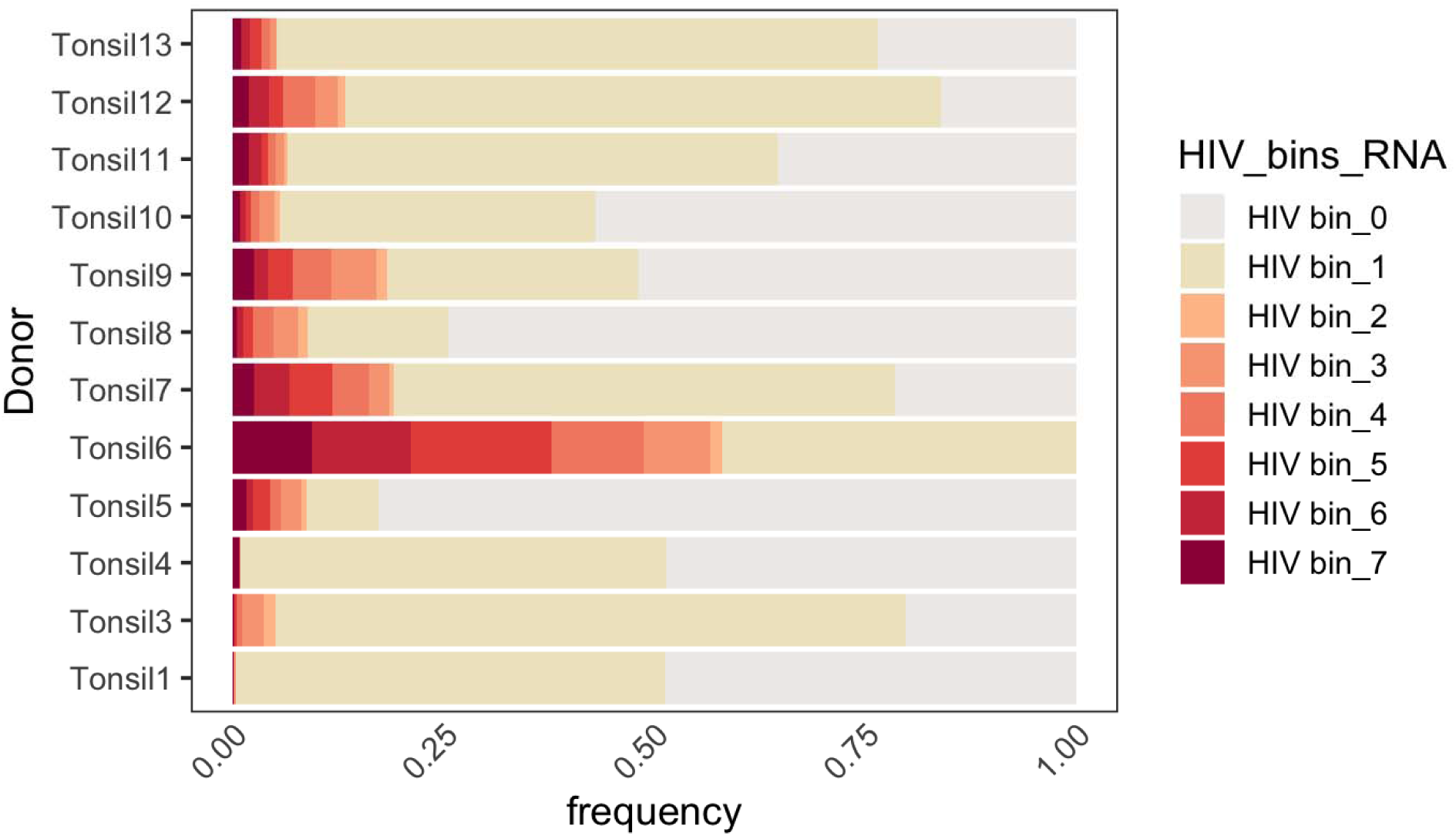
Distribution of HIV transcriptional tiers across individual donors. Stacked bar plots show the fraction of cells assigned to each transcriptional tier within individual donors. Although the relative abundance of specific tiers varied among specimens, all donors exhibited the overall continuum structure spanning exposed HIV RNA-negative and HIV RNA-positive transcriptional tiers.

**Supplementary Table 1.** Flow cytometry antibody and reporter panel used for tonsil explant phenotyping.

|  |  |  |  |  |  |
| --- | --- | --- | --- | --- | --- |
| **Target** | **Fluorochrome** | **Laser** | **Filter** | **Clone** | **Dilution** |
| CD45 | eFluor 506 | Violet405 | 525/50 | HI30 | 1:50 |
| CD11b | Super Bright 600 | Violet405 | 610/20 | M1/70 | 1:50 |
| CD8 | StarBright 670 | Violet405 | 660/20 | LT8 | 1:50 |
| CD19 | Super Bright 702 | Violet405 | 710/50 | HIB19 | 1:50 |
| Open Channel |  | Violet405 | 780/60 |  |  |
| HIV Reporter<br>(JR-FL GFP) | GFP | Blue488 | 530/30 |  |  |
| CD16 | PerCP-Cy5.5 | Blue488 | 695/40 | 3G8 | 1:50 |
| Open Channel |  | Yellow561 | 585/16 |  |  |
| HIV Reporter<br>(NL4-3 mCherry) | mCherry | Yellow561 | 610/20 |  |  |
| CD14 | PECy7 | Yellow561 | 780/60 | 61D3 | 1:50 |
| CD3 | APC | Red640 | 670/14 | UCHT1 | 1:50 |
| Viability Dye | Live-Dead Fixable<br>Near IR | Red640 | 780/60 |  | 1:1000 |
| Lineage markers, viability dye, and reporter channels for JRFL GFP and NL4-3 mCherry (NL-CI).<br>Excitation lasers, emission filters, antibody clones, and working dilutions. Acquired on Attune NxT. |  |  |  |  |  |

**Supplementary Table 2.**

| **A. Planned post-hoc comparisons from linear mixed-effects models of productive HIV infection (mCherry+ frequency, Day 8 comparison)** |  |  |  |  |  |  |
| --- | --- | --- | --- | --- | --- | --- |
| **Comparison** | **Delta**<br>**(percentage)** | **SE** | **df** | **95% CI**<br>**(lower,** | **t** | **p (Holm)** |

|  | points) |  |  | upper) |  |  |
| --- | --- | --- | --- | --- | --- | --- |
| NL4-3<br>Infected vs.<br>Uninfected | +2.237 | 0.208 | 170 | +1.692,<br>+2.781 | +10.707 | <0.0001 |
| NL4-3<br>Infected<br>+ART vs.<br>NL4-3<br>Infected | -2.022 | 0.214 | 170 | -2.582, -<br>1.463 | -9.416 | <0.0001 |

| Comparison | Delta<br>(percentage<br>points) | SE | df | 95% CI<br>(lower,<br>upper) | t | p (Holm) |
| --- | --- | --- | --- | --- | --- | --- |
| NL4-3 Infected vs.<br>Uninfected | -3.712 | 0.759 | 144 | -5.69,<br>-1.73 | -4.88 | <0.0001 |
| JR-FL Infected vs.<br>Uninfected | -1.640 | 0.788 | 144 | -3.69,<br>+0.41 | -2.08 | 0.0783 |
| NL4-3 Infected<br>+ART vs. NL4-3<br>Infected | +3.975 | 0.759 | 144 | +1.99,<br>+5.95 | +5.23 | <0.0001 |
| JR-FL Infected<br>+ART vs. JR-FL<br>Infected | +4.073 | 0.848 | 144 | +1.86,<br>+6.28 | +4.80 | <0.0001 |
| Uninfected +ART<br>vs. Uninfected | -0.002 | 0.788 | 144 | -2.06,<br>+2.05 | -0.003 | 0.9974 |

Estimated marginal mean comparisons from linear mixed-effects models testing condition effects on A) mCherry+ frequency or B) CD4+ frequency, shown as percentage-point differences (Delta) with standard errors, Kenward-Roger degrees of freedom, 95% confidence intervals, t statistics, and Holm-adjusted p-values. value ~ Condition \* Day + (1|Donor).

**Supplementary Table 3.** Cell-type composition of the integrated human tonsil single-cell RNA-seq dataset.

|  |  |  |
| --- | --- | --- |
| **Cell Category** | **Count** | **% of Total** |
| CD4 T cell | 45,075 | 31.10% |
| Memory B cell (MBC) | 21,298 | 14.70% |
| Naive CD4 T cell | 21,624 | 14.90% |
| Naive B cell (NBC) | 18,516 | 12.80% |
| CD8 T cell | 13,828 | 9.50% |
| Naive CD8 T cell | 9,607 | 6.60% |
| Cycling T cell | 7,885 | 5.40% |
| Activated NBC | 5,406 | 3.70% |
| Germinal Center B cell | 1,536 | 1.10% |
| Plasma cell | 1,521 | 1.00% |
| Innate lymphoid cell | 1,730 | 1.20% |
| Natural killer cell | 1,081 | 0.70% |
| Epithelial cell | 987 | 0.70% |
| Double-negative T cell | 695 | 0.50% |
| Monocyte/Macrophage | 486 | 0.30% |
| Dendritic cell | 294 | 0.20% |
| Unassigned | 1,227 | 0.80% |
| Granulocyte | 54 | 0.04% |
| Follicular dendritic cell | 14 | 0.01% |
| Annotated cell categories, total cell counts, and proportions of the full dataset after integration and manual curation. The dataset was dominated by lymphoid populations, with CD4 T cells comprising the largest compartment (31.1%), followed by naive CD4 T cells (14.9%), memory B cells (14.7%), naive B cells (12.8%), and CD8 T cells (9.5%). |  |  |

**Supplementary Table 4.** Distribution of cells along HIV transcriptional tiers under untreated and ART-treated conditions.

|  |  |  |  |  |  |  |
| --- | --- | --- | --- | --- | --- | --- |
| **Tier** | **HIV min** | **HIV max** | **Untreated** | **Treated** | **Total** | **Percent Treated** |
| **7** | **4.66** | **8.66** | **784** | **10** | **794** | **1.26%** |
| **6** | **2.74** | **4.65** | **782** | **14** | **796** | **1.76%** |
| **5** | **1.61** | **2.74** | **1110** | **65** | **1175** | **5.53%** |
| **4** | **0.97** | **1.61** | **948** | **164** | **1112** | **14.75%** |
| **3** | **0.38** | **0.97** | **876** | **210** | **1086** | **19.34%** |
| **2** | **0** | **0.38** | **195** | **89** | **284** | **31.34%** |
| **1** | **0** | **0** | **13862** | **6045** | **19907** | **30.37%** |
| **0** | **0** | **0** | **13002** | **4270** | **17272** | **24.72%** |
|  |  |  | **31559** | **10867** | **42,426** | **25.61%** |
| Each tier represents a discrete range of viral transcript abundance, minimum and maximum values of HIV-NL-CI log-normalized expression. Counts of untreated, ART-treated, and total T cells. Percent ART-treated cells belonging to each tier. |  |  |  |  |  |  |

